# Immunoregulatory effect of metformin in monocytes exposed to SARS-CoV-2 spike protein subunit 1

**DOI:** 10.1101/2025.09.12.675877

**Authors:** Rafael Moura Maurmann, Kierstin Davis, Negin Mosalmanzadeh, Brenda Landvoigt Schmitt, Brandt D. Pence

**Author notes:** **Corresponding Author** Brandt D. Pence, College of Health Sciences, University of Memphis, Memphis, Tennessee, TN 38152, USA.

## Abstract

**Background:** Severe COVID-19 is characterized by a hyperinflammatory state associated with an exacerbated inflammatory activation of monocytes and macrophages in the respiratory tract. Metformin has been identified as a potent monocyte inflammatory suppressor, and it has been demonstrated to attenuate inflammation in COVID-19. The mechanisms underlying metformin anti-inflammatory effects are, however, unclear. We thus sought to investigate metformin’s main interactions and their respective isolated effects in modulating monocyte inflammatory response to SARS-CoV-2 stimulation.

**Methods:** Classical human monocytes were isolated from healthy 18-40-year-old individuals and stimulated *in vitro* with recombinant spike protein subunit 1 (rS1) to assess glycolytic and oxidative metabolic responses by Seahorse extracellular flux analysis, and inflammatory gene expression by qPCR. Stimulated monocytes were either pre-treated with metformin, rotenone, S1QEL, or A769662.

**Results:** Monocytes stimulated *in vitro* with rS1 showed an increased glycolytic response associated with production of pro-inflammatory cytokines. Metformin pre-treatment reduced glycolytic activation while partially suppressing inflammation. Rotenone-dependent mitochondrial complex I inhibition was not able to replicate the same effect, and neither complex I specific ROS scavenging. Conversely, A769662 induced AMPK activation led to suppressed glycolytic inflammatory response and cytokine expression pattern similar to metformin, thus suggesting AMPK modulation as a possible central component for metformin’s mode of action upon S1 stimulation.

**Conclusions:** In summary, further investigation into the interactions underlying AMPK activity on monocytes in the context of SARS-CoV-2 may provide a better elucidation of metformin’s anti-inflammatory effect.

## Introduction

Severe acute respiratory syndrome coronavirus-2 (SARS-CoV-2) is the etiologic agent for coronavirus disease 2019 (COVID-19), a disease which has infected more than 770 million people and caused over 7 million deaths worldwide since it first outbreak by the end of 2019.^1^ While the implementation of widespread vaccination programs has considerably restrained SARS-CoV-2 transmission,^2^ the constant pathogen evolution observed during the pandemic raises serious concerns for reinfections and new outbreaks.^3^ This is a particular concern for older adults and those with comorbidities (*e*.*g*. diabetes, obesity, cardiovascular diseases) given their higher susceptibility to SARS-CoV-2 reinfections and disease severity.^4,5^ Therefore, pharmacological strategies to counteract disease severity and decrease mortality are still needed.

Hyperinflammation is a critical hallmark of severe SARS-CoV-2 infection^6^ associated with poor prognosis and increased mortality risk.^7,8^ Despite inflammatory pattern heterogeneity among individuals,^7,9^ it resembles the patterns observed in cytokine release syndromes, such as macrophage activation syndrome, thus suggesting a direct connection between dysregulated mononuclear phagocyte activation and COVID-19 severity.^10,11^ In fact, increased monocyte and monocyte-derived macrophage infiltration into the lungs of individuals with severe COVID-19 has been demonstrated by single cell RNA sequencing studies^12,13^ and postmortem analyses.^14,15^ Accordingly, a strong inflammatory activation has been detected in these cells in the lungs of critical patients.^12,16^ Markers of inflammatory activation have also been observed in circulating monocytes, denoted by high expression of cytokines, expansion of CD14+CD16+ subsets, and decreased HLA-DR expression.^7,17–19^ Altogether, these alterations on the innate immune compartment highlight the striking role of these cells in disease progression.

Therapeutic approaches targeting hyperinflammation have been widely explored for reducing COVID-19 severity and mortality.^20^ Among them, metformin, a commonly used anti-diabetic drug, has been shown to decrease mortality^21,22^ and systemic inflammation^23,24^ in COVID-19 patients undergoing treatment prior to diagnosis. Metformin has also been pointed out as a treatment for non-COVID-19 acute respiratory distress syndromes^25^ and as potent suppressor of monocyte and macrophage inflammatory activation.^24,26–31^ The mechanisms underlying metformin immunomodulatory properties are, however, still debatable, and have been mostly attributed to interference with metabolic rewiring of immune cells.^32^ Of note, metformin’s mode of action is directly related to inhibition of mitochondrial respiration through complex I blockage and consequent stimulation of AMPK due to a drop in ATP production.^33^ While some recent studies have correlated metformin anti-inflammatory properties with inhibition of complex I activity and associated ROS production independent of AMPK activity,^24,26,27^ others have pointed out to the central role of this energy sensor.^28–31^ Nonetheless, which exact mechanisms are at play in the context of COVID-19 remains an open question.

We have previously demonstrated that metformin is able to oppose hypercytokinemia in primary human monocytes stimulated with SARS-CoV-2 recombinant spike protein and live virus.^34^ Metformin reduced cytokine production and strongly inhibited both glycolytic activation and mitochondrial respiration *in vitro*. However, we did not determine the mechanisms underlying such adaptations. Therefore, in order to further investigate *in vitro* modulation of metformin observed in our previous study, we sought to isolate metformin’s main interactions, *i*.*e*. complex I inhibition and AMPK stimulation, and evaluate their impact on inflammatory response to SARS-CoV-2 stimulation.

## Methods

### Subjects

17 healthy 18-40-year-old individuals were recruited without respect to sex or race. Subjects reported to the laboratory every two weeks, and 8 mL of blood were collected into EDTA-treated vacutainer tubes by venipuncture in each visit. Blood samples were immediately used for monocyte isolation as described below.

### Monocyte Isolation

Human classical monocytes (CD14^+^CD16^-^) were isolated using immunomagnetic negative sorting kit (EasySep Direct Human Monocyte Isolation Kit, StemCell Technologies, Cambridge, MA). As we have previously described,^35^ this procedure yields a highly pure (> 85%) population of classical monocytes with depletion of intermediate (CD14^+^CD16^+^) and non-classical monocytes (CD14^-^CD16^+^) due to the presence of an anti-CD16 antibody in the cocktail. Isolated cells were diluted 10× for counting using a hemocytometer chamber. All downstream assays were performed with freshly isolated monocytes.

### Media and Reagents

All assays were performed using Seahorse XF base DMEM medium (Agilent, Santa Clara, CA) supplemented with 10 mM glucose and 2 mM L-glutamine (Millipore Sigma, St. Louis, MO). Recombinant spike protein subunit 1 (rS1) was purchased from RayBiotech (Peachtree Corners, GA). Metformin, S1QEL, rotenone, and A769662 were purchased from Millipore Sigma (St. Louis, MO).

### Seahorse Extracellular Flux Assay

Glycolysis and oxidative phosphorylation were quantified via kinetic monitoring of extracellular acidification rate (ECAR) and oxygen consumption rate (OCR), respectively, on a Seahorse XFp analyzer (Agilent, Santa Clara, CA). For all assays, monocytes were plated at 1.5x10^5^ cells per well (B-G). Wells A and H were used for background measurement without cells. All analyses were run in duplicate.

For metformin, S1QEL, rotenone, and A769662 seahorse assays, cells were incubated with media (wells B-E), or either 5 mM metformin, 5 µM S1QEL, 5 µM rotenone, or 100 µM A769662 (wells F-G) for 1 hour at 37°C in a non-CO_2_ incubator prior to Seahorse assays. 5 basal measurements were performed, followed by injection of media (wells B-C) or 300 nM spike protein (wells D-G) for a final concentration of 0 nM (wells B-C) or 30 nM spike protein (wells D-G). ECAR and OCR were then monitored serially for 30 measurements. Following the assay, images from all wells were taken at 10× magnification for cell counting in order to adjust raw measurements for cell number. Cell culture supernatants were then removed, pooled by duplicate, and stored at -80°C. Cells were then lysed with 100 μl Trizol (Thermo Fisher Scientific, Waltham, MA), pooled by duplicate, and stored at -80°C.

### Gene Expression

RNA isolation was performed using the Trizol method based on manufacturer’s instructions for cell lysates retrieved from Seashore assays. Isolated RNA was quantified on Nanodrop Lite instrument (Thermo Fisher Scientific). 200-600 ng of RNA was reverse-transcribed to cDNA using a High-Capacity cDNA Reverse Transcription Kit (Thermo Fisher Scientific, Waltham, MA). Gene expression was analyzed using commercial pre-validated gene expression assays and Taqman reagents (Thermo Fisher Scientific, Waltham, MA) on QuantStudio 6 instrument (Thermo Fisher Scientific). Relative gene expression was quantified using the 2^-ΔΔCt^ method^36^ against ACTB as housekeeping gene. Primer/probe IDs were: ACTB (Hs03023943_g1); IL1B (Hs01555410_m1); IL6 (Hs00174131_m1); CXCL8 (Hs00174103_m1); TNF (Hs00174128_m1); IL10 (Hs00961622_m1).

### Data Processing and Statistical Analysis

All data processing and statistical analyses were performed in R v.4.3.1.^37^ Isolated monocytes from each individual were subjected to all treatments for each experiment, so data were paired and analyzed using within-subjects designs. Data were checked for normality using Shapiro-Wilk test and analyzed by one-way repeated measures ANOVA (RM-ANOVA, for data which met the normality assumption) or Friedman’s test (for data which did not meet the normality assumption). For analyses with significant main effects, *post hoc* mean separation was performed using pairwise paired T tests (for RM-ANOVA) or pairwise Wilcoxon signed-rank tests (for Friedman’s tests) with p-value adjustment using the Holm-Bonferroni method.^38^ Significance cutoff was p<0.05. For cluster analysis cytokine expression data of all treatment was combined, standardized, and modeled using K-means approach.

All raw data and analytical scripts (as R markdown files) are available in a dedicated FigShare repository.^39^

### Ethical Approval

All human subject activities were approved by the Institutional Review Board at the University of Memphis under protocol PRO-FY2021-476, and subjects provided written informed consent prior to enrollment.

## Results

### Metformin Suppresses Inflammatory Response to Spike Protein

SARS-CoV-2 spike protein has been previously reported to trigger inflammatory response in myeloid cells.^40,41^ We have previously demonstrated metformin treatment to suppress inflammatory activation upon spike protein stimulation.^34^ Pre-treatment with metformin reduced glycolytic response to rS1 (**Figure 1A**) and strongly inhibited mitochondria respiration (**Figure 1B**) in Seahorse assays. This metabolic response was accompanied by the suppression of IL6 cytokine gene expression, as well as a reduction in IL10 expression (**Figure 1C**). TNF expression was partially reduced but did not reach statistical significance due to limited statistical power (**Figure 1C**). No significative changes were observed regarding IL1B expression (**Figure 1C**).

**Figure 1.**
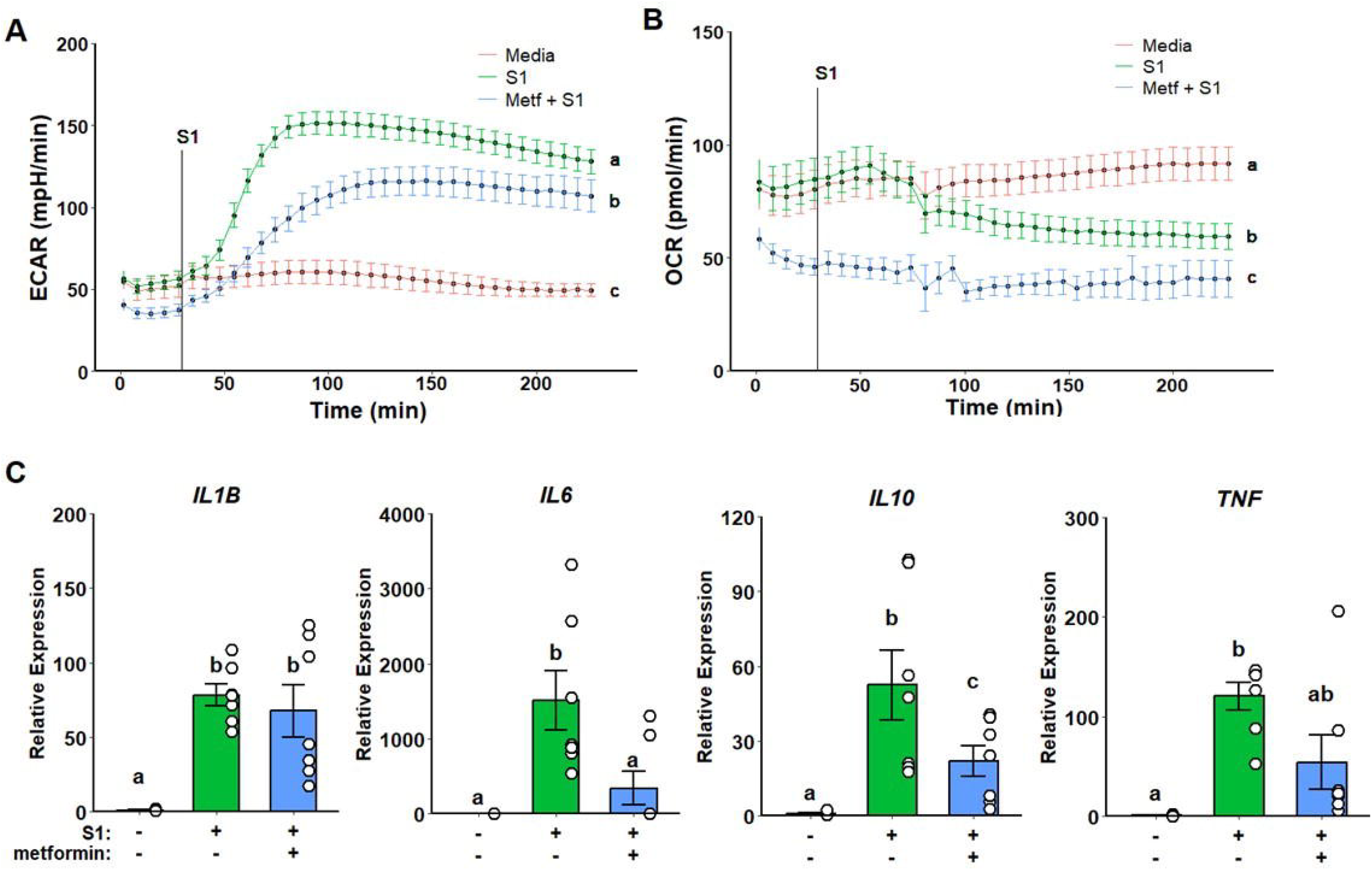
Metformin attenuates glycolytic activation and suppresses mitochondrial respiration while partially decreasing inflammatory response to SARS-CoV-2 spike protein subunit 1 (rS1). Metformin reduces extracellular acidification rate (ECAR) in response rS1 (A) and ablates oxygen consumption rate (OCR) (B). Additionally, metformin alleviates upregulation of inflammatory cytokine gene expression (C).

### Metformin Inflammatory Suppression is not Dependent on Complex I Inhibition and ROS Production

To test whether metformin inflammatory suppression is dependent on its modulation of complex I activity, we first pre-treated monocytes with rotenone, a complex I inhibitor. While no changes were observed in glycolytic response (**Figure 2A**), mitochondrial respiration was strongly suppressed (**Figure 2B**). However, no effects on cytokine expression were observed following rotenone pre-treatment when compared to spike treatment alone (**Figure 2C**).

**Figure 2.**
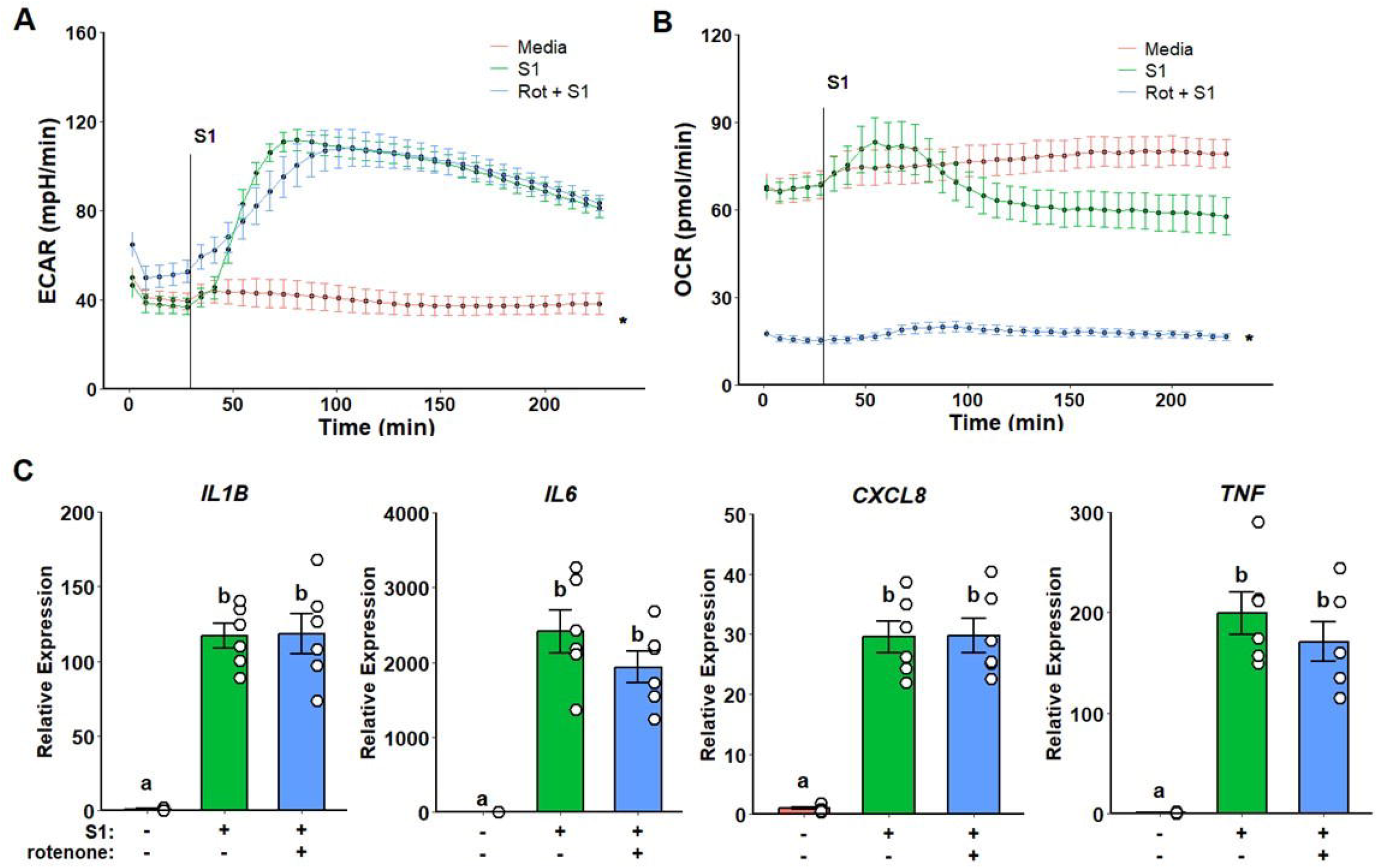
Rotenone suppresses mitochondrial respiration but not glycolytic activation or inflammatory response to SARS-CoV-2 spike protein subunit 1 (rS1). Extracellular acidification rate (ECAR) response is unaltered upon rotenone pre-treatment (A) while oxygen consumption rate (OCR) is completely suppressed. No differences were observed regarding inflammatory gene expression (C).

Next, we choose to investigate whether metformin inhibition of ROS production by complex I instead of complex I activity *per se* modulates inflammatory response. Cells were pre-treated with S1QEL, a complex I superoxide-specific scavenger. No differences in glycolytic response and mitochondrial respiration were observed (**Figure 3A-B**). Likewise, no differences were detected in gene expression when compared to spike treatment alone (**Figure 3C**).

**Figure 3.**
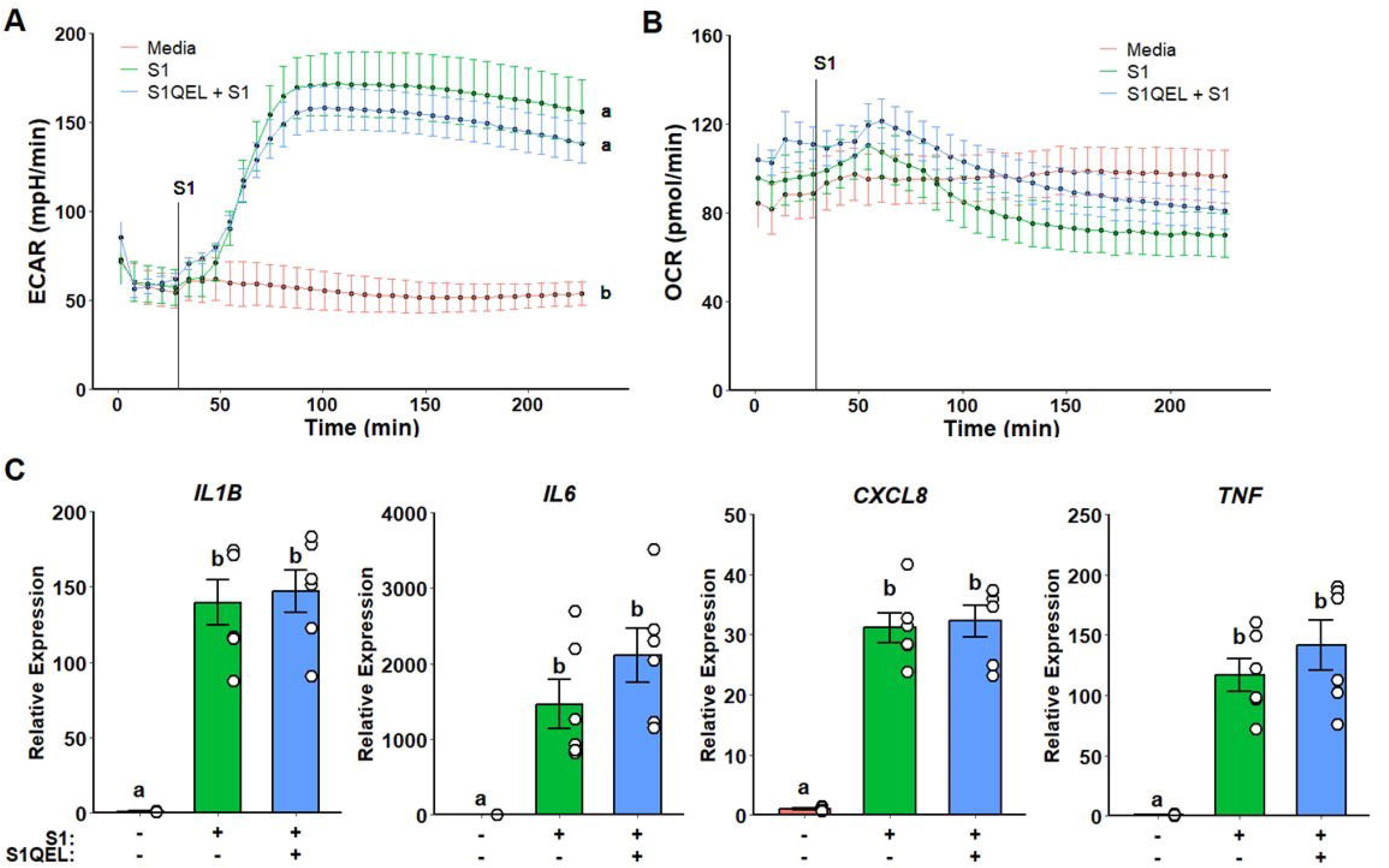
Complex I reactive oxygen species scavenging does not alter metabolic or inflammatory response to SARS-CoV-2 spike protein subunit 1 (rS1). Unaltered extracellular acidification rate (ECAR) (A) and oxygen consumption rate (OCR) (B) responses on S1QEL pre-treated cells upon rS1 stimulation. No differences were observed regarding inflammatory gene expression (C).

### Metformin Inflammatory Modulation is Independent of AMPK Activation

Given that metformin-related effects on complex I did not evoke the previously observed inflammatory effects, we assessed whether metformin action was dependent on AMPK activation. Monocytes pre-treated with A769662, a potent specific AMPK activator, displayed a suppressed glycolytic response (**Figure 4A**) without any effects on mitochondrial respiration (**Figure 4B**). Furthermore, no significative differences were detected in cytokine gene expression (**Figure 4C**).

**Figure 4.**
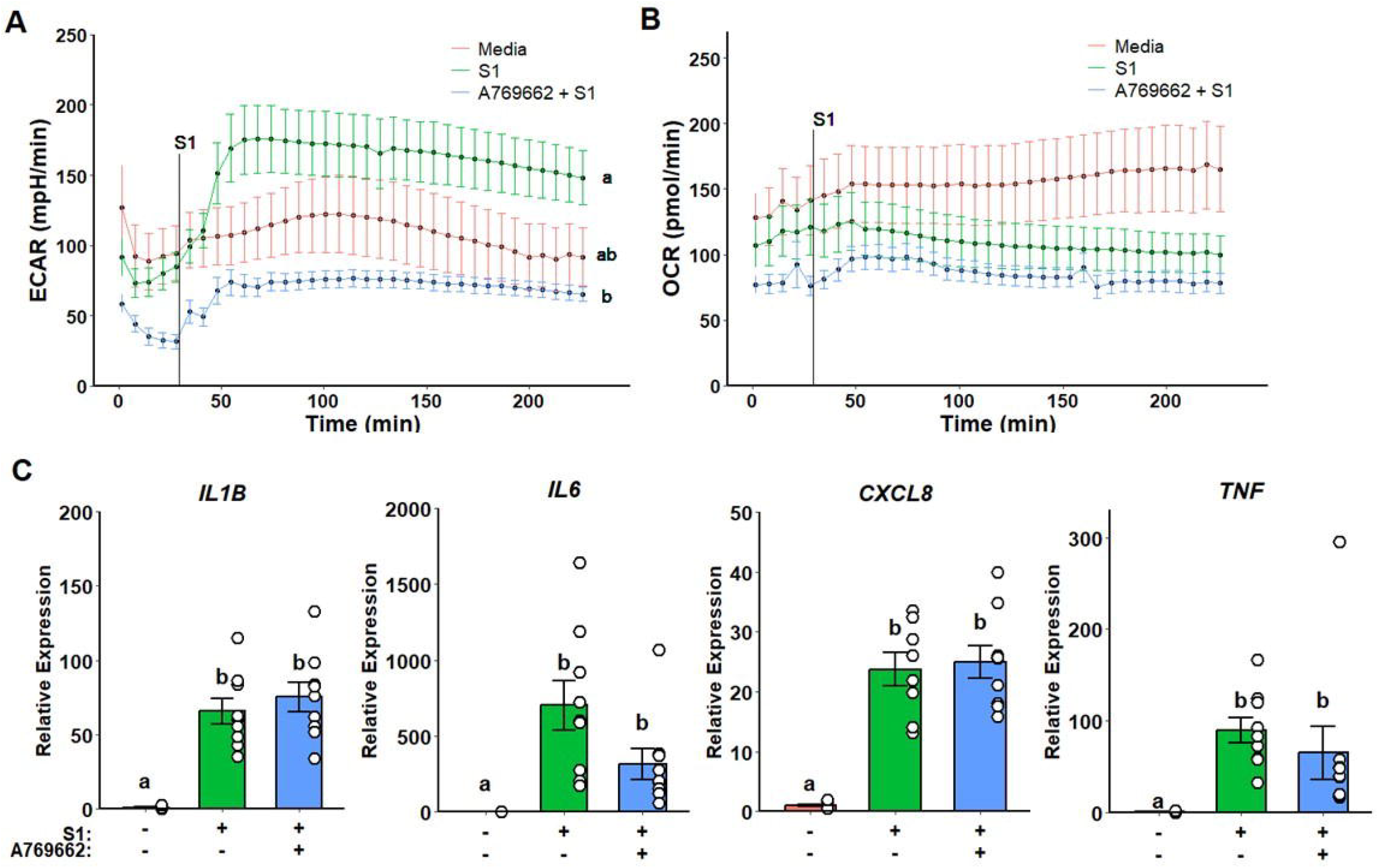
AMPK activation ablates glycolytic response but does not alter mitochondrial respiration and inflammatory response to SARS-CoV-2 spike protein subunit 1 (rS1). A769662 suppresses extracellular acidification rate (ECAR) in response rS1 (A) without altering oxygen consumption rate (OCR) (B). No significant differences were observed regarding inflammatory gene expression (C).

### Metformin Treatment and AMPK Activation Display Similar Inflammatory Profiles

Despite non-significative differences being detected on the inflammatory profile of cells pre-treated with AMPK activator, it resembles the pattern observed on metformin pre-treatment. We thus performed a cluster analysis to evaluate the similarity between the inflammatory profile outputs (IL-1B + IL6 + TNF) of each treatment. As expected, metformin and A769662 treatments where clustered together alongside non-stimulated monocytes, while rotenone and S1QEL treated monocytes clustered with most of media treated wells stimulated with spike protein (**Figure 5**).

**Figure 5.**
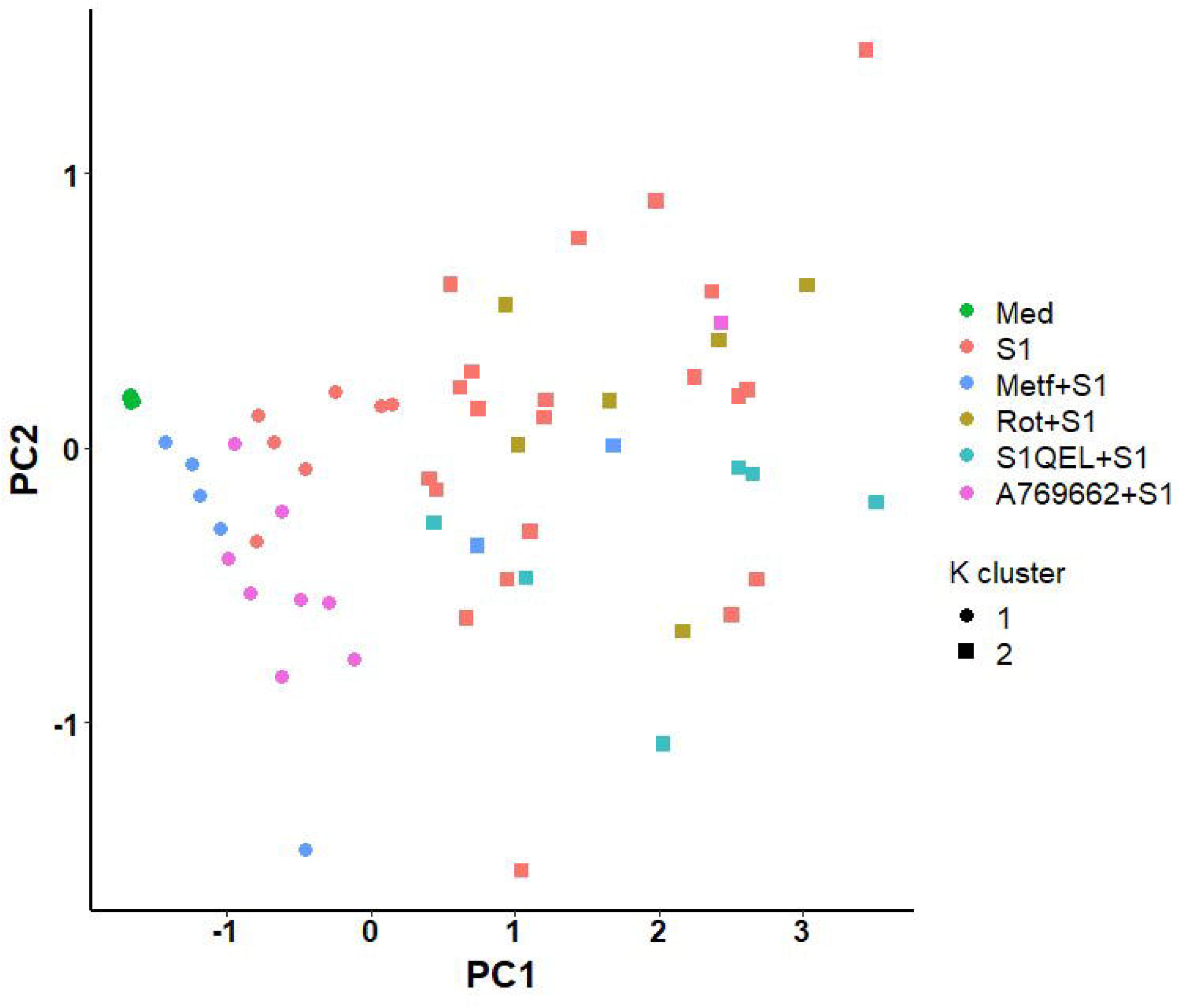
Metformin and AMPK activation display a similar inflammatory profile in response to SARS-CoV-2 spike protein subunit 1 (rS1). Metformin and A769662 treated monocytes were clustered together with non-stimulated monocytes, while rotenone and S1QEL treated cells were clustered with most of media treated wells stimulated with spike protein.

## Discussion

We have previously demonstrated metformin abrogates monocyte inflammatory response to SARS-CoV-2 recombinant spike protein and live virus *in vitro*.^34^ While we observed similar results in the present study, we detected a milder suppression regarding to cytokine expression when replicating the experiment. Nevertheless, metformin was still able to suppress glycolytic inflammatory activation and mitochondrial respiration. To elucidate the possible mechanisms underlying metformin related adaptations, we sought to isolate metformin’s main interactions. We first attempted to inhibit complex I with rotenone, but no effects were observed in terms of inflammatory response. Similarly, directly inhibiting ROS production on complex I using S1QEL did not elicit any significant effect. Therefore, we attempted to evaluate the indirect effect of metformin on AMPK. Similar to metformin, direct AMPK activation led to suppressed glycolytic inflammatory response, with a low non-significant impact on cytokine expression. However, despite the lack of significant changes in gene expression, AMPK activation elicited a cytokine expression pattern comparable to metformin, thus suggesting AMPK modulation as a possible central node for metformin’s mode of action upon S1 stimulation.

Despite the known anti-inflammatory properties of metformin, its mode of action remains controversial, especially regarding its dependency on AMPK modulation.^27,32^ For instance, Soberanes et al.^26^ demonstrated metformin to attenuate particulate matter-induced IL-6 release from alveolar macrophages by inhibiting mitochondrial complex I and consequently suppressing ROS signaling and the opening of Ca^2+^ release-activated Ca^2+^ channels, central nodes for IL-6 generation. In accordance, Kelly et al.^27^ also showed blunted LPS-induced IL-1β expression in murine bone marrow-derived macrophages due to mitochondrial complex I activity and ROS production inhibition by metformin treatment. It is important to note that in both studies metformin anti-inflammatory effects were independent of AMPK activity and mostly attributable to ablation of ROS generation. While we did not measure mitochondrial ROS production, rotenone is known to potentiate its generation,^42^ which could thus account for the unaltered inflammatory response observed in the present study. However, in Kelly’s et al.^27^ work rotenone treatment was not only able to replicate the same effect as metformin but also promoted a reduction in ROS production. Conversely, a recent study demonstrated metformin’s effect to be also independent of inhibition of mitochondrial ROS production, as pharmacological ROS stimulation was unable to restore inflammatory response to LPS following metformin treatment.^24^ The authors attributed metformin’s effect to limited generation and oxidation of mitochondrial DNA, therefore decreasing NLRP3 inflammasome signaling. In this context, combining rotenone to S1QEL treatment may hold a further elucidation whether inhibiting mitochondrial complex I depends on ROS inhibition to suppress rS1 inflammatory stimulation. Regardless of ROS production, those studies highlight the inhibiting mitochondrial complex I as a major node for metformin’s mode of action, which we failed to demonstrate.

One of the mainstream effects attributed to metformin is its indirect stimulation of AMPK through changes in cell energy balance.^33^ In fact, this phenomenon is associated with most of the benefits of metformin treatment in a plethora of cell types.^42,43^ AMPK activation antagonizes metabolic inflammatory response by limiting glycolytic capacity and driving oxidation of substrates in mitochondria.^44^ This metabolic rewiring interference has been correlated with promotion of inflammatory to anti-inflammatory macrophage switch in a plethora of pathological contexts such as obesity,^30^ ischemic stroke,^45^ wound healing,^46^ and cancer.^47^ However, a precise definition of the mechanisms underlying metformin AMPK-dependent effect remains uncovered as it has been attributed to the modulation of distinct inflammatory axis.^28–31^ Of note, it has been shown to indirectly downregulate NF-κB transcriptional activity consequently suppressing TNF-α expression and ROS production,^31^ while other studies have attributed it to NLRP3 modulation and JKN1 phosphorylation.^28,29^ Nonetheless, we did not observe a significative effect on cytokine expression following AMPK stimulation despite suppressed inflammatory glycolytic activation. Although the evidence provided in this study is not strong enough to demonstrate that metformin acts through AMPK to modulate cytokine production, we can still draw a connection between those based on their similar cytokine profile.

An important limitation of this study is that we restricted inflammatory response evaluation to metabolic measures and cytokine expression. For instance, Xian et al.^24^ demonstrated a very limited effect of metformin on cytokine expression upon LPS stimulation but with a pronounced modulation of NLRP3 inflammasome. Besides, we did not include ROS assessment in our analysis, which, despite the controversy on ROS involvement, may provide additional clues of relevant mechanisms to be explored in our model. Additionally we included a small sample size, which may blunt demonstrating some of the interrogated mechanisms due to the large effect size. Nevertheless, we include a healthy young population which we believe excludes extra variance from comorbidities such as aging and provides a better mechanistic evaluation.

## Conclusions

A precise definition of metformin’s mode of action is a challenging task given its connection with pleiotropic metabolic hubs, thus leading to controversial findings. While the tested effects of individualized key metformin’s interactions did not elicit the same response, we were able to identify AMPK activation as a promising central player on our model. We thus believe that a further investigation into the interactions underlying AMPK activity on monocytes stimulated with S1 may hold a potential area of interest.

## Acknowledgments

The authors would like to acknowledge the participants in this study.

## Author Contributions

BP conceived the study. BP designed experiments. RMM, KD, NM, BLS, and BP collected data. RMM and BP analyzed data. RMM prepared the first manuscript draft. BP edited the manuscript draft. All authors read and approved the final manuscript.

